# P681 mutations within the polybasic motif of spike dictate fusogenicity and syncytia formation of SARS CoV-2 variants

**DOI:** 10.1101/2022.04.26.489630

**Authors:** Alona Kuzmina, Nofar Atari, Aner Ottolenghi, Dina Korovin, Ido Cohen lass, Benyamin Rosental, Elli Rosenberg, Michal Mandelboim, Ran Taube

## Abstract

The rapid spread and dominance of the Omicron SARS-CoV-2 over its Delta variant has posed severe global challenges. While extensive research on the role of the Receptor Binding Domain on viral infectivity and vaccine sensitivity has been documented, the role of the spike _681_PRRAR/SV_687_ polybasic motif is less clear. Here we monitored infectivity and vaccine sensitivity of Omicron SARS-CoV-2 pseudovirus against sera samples that were drawn four months post administration of the third dose of BNT162b2 mRNA vaccine. Our findings show that relative to Wuhan-Hu and Delta SARS-CoV-2, Omicron displayed enhanced infectivity and a sharp decline in its sensitivity to vaccine-induced neutralizing antibodies. Furthermore, while the spike proteins form Wuhan-Hu (P681), Omicron (H681) and BA.2 (H681) pseudoviruses modestly promoted cell fusion and syncytia formation, Delta spike (P681R) displayed enhanced fusogenic activity and syncytia formation capability. Live-viruses plaque formation assays confirmed these findings and demonstrated that relatively to the Wuhan-Hu and Omicron SARS-CoV-2, Delta formed more plaques that were smaller in size. Introducing a single P681R point mutation within the Wuhan-Hu spike, or H681R within Omicron spike, restored fusion potential to similar levels observed for Delta spike. Conversely, a R681P point mutation within Delta spike efficiency abolished fusion potential. We conclude that over time, the efficiency of the third dose of the Pfizer vaccine against SARS CoV-2 is waned, and cannot neutralize Omicron. We further verify that the P681 position of the viral spike dictates fusogenicity and syncytia formation.

## Introduction

In late 2019, an emergent betacorinavirus, Severe Acute Respiratory Syndrome Coronavirus 2 (SARS-CoV-2) was identified in China and defined as the cause for severe respiratory COVID-19 disease in humans. Two years following this outbreak, it is clear that the pandemic is here to stay, leading to about 4% mortality and causing global effects on economic and quality of life [1, 2].

Efforts to minimize spread of the virus led to major efforts to develop and authorize two mRNA vaccines, Pfizer (BNT162b2) and Moderna (mRNA-1273) that target the receptor binding domain (RBD) of the original Wuhan-Hu viral spike. These efforts have matured to a successful vaccine that provided efficient protection against viral infection and disease development [3-8] [9-13] [10, 13, 14]. However, rapid spread and viral evolution led to the emergence of new viral variants that partly resist the vaccines, as they exhibit improved affinity to the ACE2 cell receptor and successful escape from neutralizing antibodies. Early major variants of concern (VOC) included Alpha [15] and Beta [16, 17] [18-28], that were then dominated by the Delta SARS-CoV-2 that led to severe clinical COVID19 symptoms [29-31]. To combat emergence of new variants and waning of vaccine-elicited antibody response, a third dose of mRNA vaccine has been approved in August 2021, first in Israel and later worldwide. This boost has shown to be very effective at inducing high neutralizing antibody protection and preventing complications of disease development and hospitalization [32]

In November 2021 another VOC has emerged in South Africa and Botswana -BA.1/B.1.1.529 also designated Omicron. This variant has since quickly become worldwide dominant. Omicron harbors up to 59 mutations throughout its genome, with 36 of them are positioned within the spike and 15 specifically within the RBD. Subsequent reports show that Omicron partially escapes from immunity elicited by infection or vaccines, with boosted individuals presenting waned vaccine efficacy over time. However, these reports also show that compared to Delta and Alpha, Omicron results in lower incidences of severe COVID19 [33-40]. To further boost the protection of vaccine-elicited protection against Omicron and its lineages, Israel has recently initiated a fourth boost that is currently being evaluated for its efficiency and indicates a short-term protection against SARS CoV-2 variants with no boosting [41-43].

While the involvement of mutations within the RBD of spike are extensively being investigated for a role in antibody-neutralization and viral infectivity, the role of _681_PRRAR/SV_687_ mutations located at the S1/S2 boundary of spike and located within the Furin Cleavage Site (FCS) is less clear. This polybasic motif is unique to SARS-CoV-2 and has been previously reported to promote viral entry in a TMPRSS2 dependent. TMPRSS2 on the target cell surface and cathepsins B and L in endosomes then cleave the S2′ site and activate the fusion machinery, depending on the relative availability of these enzymes [44, 45]. The polybasic region in spike has also recently shown to enhance fusion activity and syncytia formation [46, 47] [48-50], and also facilitate TMPRSS2 endocytic viral entry that allows the evasion of IFITM2 innate immune response [51]. Finally, Omicron has been also reported to exhibit lower replication levels in lung and gut cells, and its spike was less efficiently cleaved compared to Delta. These differences were mapped to the entry efficiency of the virus to specific cell types and effectively correlated with higher cellular TMPRSS2 RNA expression. Overall, Omicron spike inefficiently used TMPRSS2, which promotes cell entry through plasma membrane fusion, with greater dependency on cell entry through the endocytic pathway. Consistent with the inefficient S1/S2 cleavage and inability to use TMPRSS2, syncytium formation by Omicron spike was impaired compared with the Delta spike. This inefficient spike cleavage of Omicron with a shift in cellular tropism away from TMPRSS2-expressing cells, with implications for altered pathogenesis [52-54].

Here we tested the neutralization potential of sera drawn from individuals who were fully vaccinated with a third dose of the Pfizer BNT162b2 vaccine - four months post dose administration - against the Wuhan-Hu SARS CoV-2 pseudovirus or its Delta and Omicron variants. We show that relative to the wild-type Wuhan-Hu SARS-CoV2 and its Delta variant, Omicron displayed a sharp decrease in neutralization sensitivity, as well as elevated infectivity levels. Live virus infection plaque assay confirmed that Delta variant formed more plaques, which were less unified in their size and overall smaller when compared to the Wuhan-Hu and Omicron SARS CoV-2. We then investigated the role of the P681 residue in the spike FCS in enhancing cell fusion and syncytia formation. We imaged fusion syncytia formation between target cells expressing Wuhan-Hu, Delta, Omicron and BA.2 spike proteins, either by a GFP-split system that monitored GFP expression as a measure of cell fusion upon spike expression [46], or by following fusion of transduced cells that express two different fluorescent proteins. While the Wuhan-Hu spike, that carries a P681, did not enhance fusogenic activity and syncytia formation, cells expressing Delta-R681 spike efficiently enhanced fusion and syncytia formation between target cells. Introducing a single P681R mutation in the spike of Wuhan-Hu-SARS-CoV-2, restored fusogenic potential and enhanced syncytia formation of this spike. Parallelly, expression of Delta-spike protein carrying a single R681P mutation, abolished fusion potential and syncytia formation. Omicron spike, that exhibits a P681H also displayed low levels of fusion potential and syncytia formation – similar to those of the Wuhan-Hu spike. Introducing an H681R in spike of Omicron, restored cell fusion potential and enhanced syncytia formation, similar to those exhibited by Delta SARS-CoV-2. We therefore conclude that the P681 position of spike is critical for mediating cell fusion and syncytia formation.

## Methods

### Virus and viral culture

SARS-CoV-2 Delta and Omicron variants were isolated from the combined nasopharyngeal throat swab of a COVID-19 patient in Israel. The viral culture was performed in a biosafety level 3 facility as we described previously. Briefly, TMPRSS2-expressing VeroE6 (VeroE6/TMPRSS2) cells (JCRB Cat^#^JCRB1819) were seeded with 100 μL of minimum essential medium (MEM) (Thermo Fisher Scientific) at 4 × 10^4^ cells in a 96-well plate and incubated at 37°C in a carbon dioxide incubator until confluence for inoculation. Each well was inoculated with 30 μL of clinical specimen. The virus-induced cytopathic effect was examined. Cultures with more than 50% virus-induced cytopathic effect were expanded to large volume, and 50% tissue culture infective dose (TCID_50_) was determined. The whole-genome sequence of the culture isolates was confirmed using nanopore sequencing as we described previously

### Fusion assays

For cell–cell fusion measurements, Vero-E6 cells stably expressing GFP1-10 and GFP11 were co-cultured at a 1:1 ratio and were transfected with a total of 1µg of phCMV-SARS-CoV2-Spike with TransIT-X2 (Mirus) in a 24-well plate. At 24 h post-transfection, GFP fluorescence images were documented using Olympus IX73. The GFP area and number of nuclei were quantified using ImageJ software. Fusion was defined as percentage of GFP pixels.

### Cells

Vero-ACE2 stable cells were cultured at 37^0^C in a 5% CO2 incubator. Cells were grown in Dulbecco’s Modified Eagle Medium (DMEM) high glucose (Gibco), supplemented with 10% fetal bovine serum (FBS), 2mM GlutaMAX (Gibco) and 100U/ml penicillin-streptomycin. Vero -ACE2 expressing cells were generated by stable transduction with lentivirus expressing human ACE2. Our pseudoviruses were standardized for equal loads by monitoring p24 levels by ELISA. *E*.*coli* DH5αbacteria were used for transformation of plasmids coding for lentivirus packaging DNA and SARS CoV-2 spike. A single colony was picked and cultured in LB broth with 50µg penicillin at 37^0^C at 200 rpm in a shaker for overnight.

### Generation of Vero-hACE2 stable cell line

hACE2 (received from S. Pohlmann lab, University Göttingen, Germany) was re-cloned into lentiviral expression vector. Lentiviral particles were produced as described previously [55] Briefly, Vero cells were stably transduced with lentivirus expressing ACE2. Cells were analyzed for hACE2 expression by FACS, using biotinylated-labeled spike (ACROBiosystems). High ACE2 expressing cells were sorted using FACS Aria. ACE2 expression was periodically monitored by FACS.

### Mutagenesis of Spike mutants

QuikChange Lightening Site-Directed Mutagenesis kit was used to generate amino acid substitutions in the pCDNA spike plasmid (received from S. Pohlmann lab, University Göttingen, Germany), following the manufacturer’s instructions (Agilent Technologies, Inc., Santa Clara, CA). For each mutant the relative oligos that harbored the required mutation were employed. Omicron spike was synthetically synthesized by IDT.

### Generation of pseudotyped lentivirus and neutralization assays

Pseudotyped viruses were generated in HEK293T cells. Briefly, LTR-PGK luciferase lentivector was transfected into cells together with other lentiviral packaging plasmids coding for Gag, Pol Tat Rev, and the corresponding wild type or mutate spike envelopes. Transfections were done in a 10cm format, as previously described and supernatant containing virus were harvested 72hr post transfection, filtered and stored at -80^0^C [55]. Neutralization assays were performed in a 96 well format, in the presence of pseudotyped viruses that were incubated with increasing dilutions of the tested sera (1:2000; 1:8000: 1;32000: 1:128000) or without sera as a control. Cell-sera were for 1hr. at 37°C, followed by transduction of HEK-ACE2 cells for additional 12 hr. 72hr post transduction, cells were harvested and analyzed for luciferase readouts according to the manufacturer protocol (Promega). Neutralization measurements were performed in triplicates using an automated Tecan liquid handler and readout were used to calculate NT_50_ – 50% inhibitory titers concentration.

### Pseudoviruses quality control and Tittering

To determine the titers of pseudoviruses, 1×10^5^ ACE2 stable HEK cells were plated in a 12-well plate. 24 h later, decreased serial dilutions of pseudovirus were used to transduce cells. 48 h post transduction, cells were harvested and analyzed for their luciferase readouts. p24 ELISA measurements were conducted to ensure equal loads.

### Expression of spike proteins and monitoring fusion potential

Vero-ACE2 cells were transfected with the indicated spike protein and fusion potential and syncytia formation were visualized by microscopy, 72hr later. In addition, we used the GFP-split system, where GFP10 and GFP11 Vero cells were equally mixed and then transfected with the indicated spike protein.

### Analysis of plaque formation

Vero E6 cells were grown in complete DMEM/10% FCS. Cells were seeded in 6 well plates at a concentration of 10^6^ cells/well with 10% FCS MEM-EAGLE medium, and incubated at 37°C. 24 hours later, the cells were infected with the following SARS-CoV-2 variants: 1. Wild Type sub lineage B.1.1.50 (hCoV-19/Israel/CVL-45526-ngs/2020), 2. Delta, B.1.617.2 (hCoV-19/Israel/CVL-12804/2021), and 3. Omicron BA.1 (hCoV-19/Israel/CVL-n49814/2021) as 10 fold serial dilutions of 100TCID50 for 1h at 33°C. The cells then were washed and MEM-EAGLE medium containing 2% FCS and 0.5% NuSieve agarose was added. The cells were then incubated at 33°C for 5 days additional days. For fixation, 10% of Trichloroacetic acid was added to each well for 10 minutes at room temperature. The gel was washed and the plates were stained with Gentian Violet which contains 4% Formaldehyde to visualize plaques.

### Image analysis

Image analysis was done using the Scikit-image library [56] in Python 3.1. Images for each channel (GFP – fused cells, and DAPI – nuclei stain) were loaded separately. Histograms for all images were equalized. Using the Otsu threshold algorithm on the GFP channel, borders of the fused cells were determined. The cell regions in each image were isolated and labeled using the scikit-image label algorithm, where each region corresponds to a fused cell. These regions were then iterated over, and used to individually mask the corresponding region over the DAPI channel. For each iteration the number of nuclei in the masked region, and therefore the fused cell, was assessed using the local maxima in the intensity of the DAPI image, with a minimum distance of 3 pixels between each peak. The pixel area of each region was also measured, giving us, in addition to nuclei number in each cell, the its area also. For statistical analysis, only regions with 2 or more nuclei were selected. Statistical analysis was done using a two-sided independent t-test from the stat module of the scipy [57] library. Graphing was done using the matplotlib and seaborn libraries [58]. Additional libraries used – pandas and numpy [59].

### Quantification and statistical analysis

Statistical analyses were performed using GraphPad Prism. Measured statistical significance was calculated between experiments by a two-tailed Student’s t test - P≤0.001. Error bars throughout all figures represent one standard deviation. Specific details on statistical tests and experimental replicates can be found in the figure legends.

## Results

### Neutralizing sensitivity and viral infectivity of Omicron SARS CoV-2 pseudovirus and its FCS mutants against post-vaccination sera

We acquired blood samples from a cohort of individuals who were fully vaccinated with the third dose of the BNT162b2 Pfizer vaccine. Sera was drawn four months post administration of the third dose (n=20). Our cohort was tested for neutralizing potential against the original Wuhan-Hu SARS CoV-2, its Delta and Omicron variants. We show that relative to the Wuhan-Hu SARS-CoV-2 pseudovirus, Omicron exhibited a x26 fold decrease in its neutralization sensitivity to the tested post-vaccinated sera (**Figure 1A**). Omicron also exhibited a x13 fold decrease in neutralization sensitivity relative to the Delta pseudovirus. We also monitored the infectivity levels of the Omicron and Delta variants relative to the wild type Wuhan-Hu pseudovirus. Omicron was x5 fold more infective than the Wuhan-Hu SARS CoV-2. Moreover, we confirmed previous data reporting a x2-3 fold increase in Delta variant infectivity relative to the wild type SARS CoV-2 pseudovirus (**Figure 1B**) [31].

**Figure 1:**
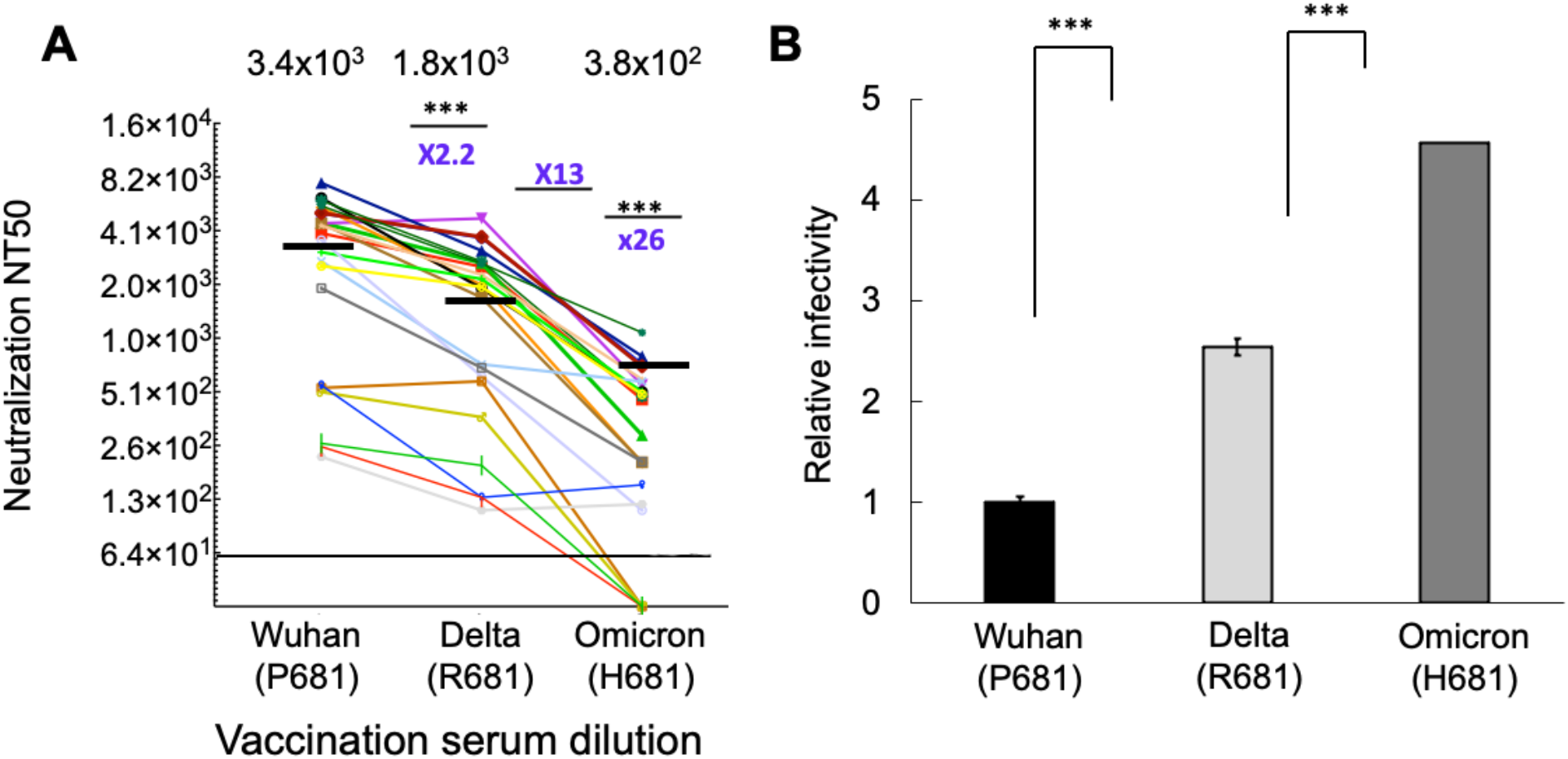
Neutralization potential against SARS CoV-2 and its recent variants of concern. **A. Neutralization sensitivity of Omicron and Delta pseudoviruses** - neutralization assays were performed with sera samples of individuals that were fully vaccinated and drawn four months post administration of the Pfizer-vaccine. For neutralization measurements, sera samples were incubated with the indicated pseudoviruses, followed by transdcution of HEK293T-ACE2 target cells. 48 hr post transduction, cells were harvested and their transduction was monitored by measuring luciferase readings. Neutralizing potency was calculated at increased serial dilutions, relative to transduced cells with no sera added. Neutralization, NT_50_ is defined as the inverse dilution that achieved 50% neutralization. Results are the average of two independent biological experiments. Triplicates were performed for each tested serum dilution. Black bars represent geometric mean of NT_50_ values, indicated at the top. Statistical significance was determined using one tailed t-test ***<0.001. **B. Infectivity levels of the Wuhan-Hu SARS CoV-2 pseudoviruses and its Delta or Omicron variants** Pseudoviruses carrying Wuhan-Hu SARS-CoV-2 spike or its Delta (P681R) and Omicron (P681H) spikes were used to transduce HEK293T-ACE2 target cells. Equal viral loads were normalized based on p24 protein levels. 48 hr post transduction, cells were harvested and their luciferase readouts were monitored. Bar graphs show mean values ± SD error bars of three independent experiments.

As our platform used single-round pseudoviruses, the term transduction is more suitable than infectivity that would imply the use of infections SARS-CoV-2. Moreover, a close correlation exists between pseudovirus and live viruses regarding the measurements of viral entry and neutralization potential.

### Fusogenic potential of wild type SARS-CoV-2 and its Delta or Omicron variants

We next expressed spike proteins from the Wuhan-Hu, Delta and Omicron in Vero-GFP-split cells and documented GFP expression as a measure of fusion potential (**Figure 2A**) [49]. The Wuhan-Hu spike carries a Proline at position 681, part of the poly-basic motif in spike. P681 is mutated to R681 within the Delta spike, or H681 in Omicron variants. Our data demonstrated that upon expression of the Delta (R681) spike in split Vero-GFP cells, the efficiently of cells to fuse and form syncytia was substantially enhanced. Delta exhibited relatively large syncytia, when compared to the other SARS-CoV-2 pseudoviruses. In contrast, spike from Wuhan-Hu, Omicron, BA.1 and BA.2 exhibited a modest fusogenic potential and their syncytia were rather small (**Figure 2B**). GFP expression as a measurement for fusion potential was also quantitated and confirmed our results. Image analysis and quantitation of nuclei per cell for each of the variants and cell area was calculated based on the expression of GFP and DAPI and was performed using the Scikit-image library [56] in Python 3.1 (**Figure 2C**).

**Figure 2:**
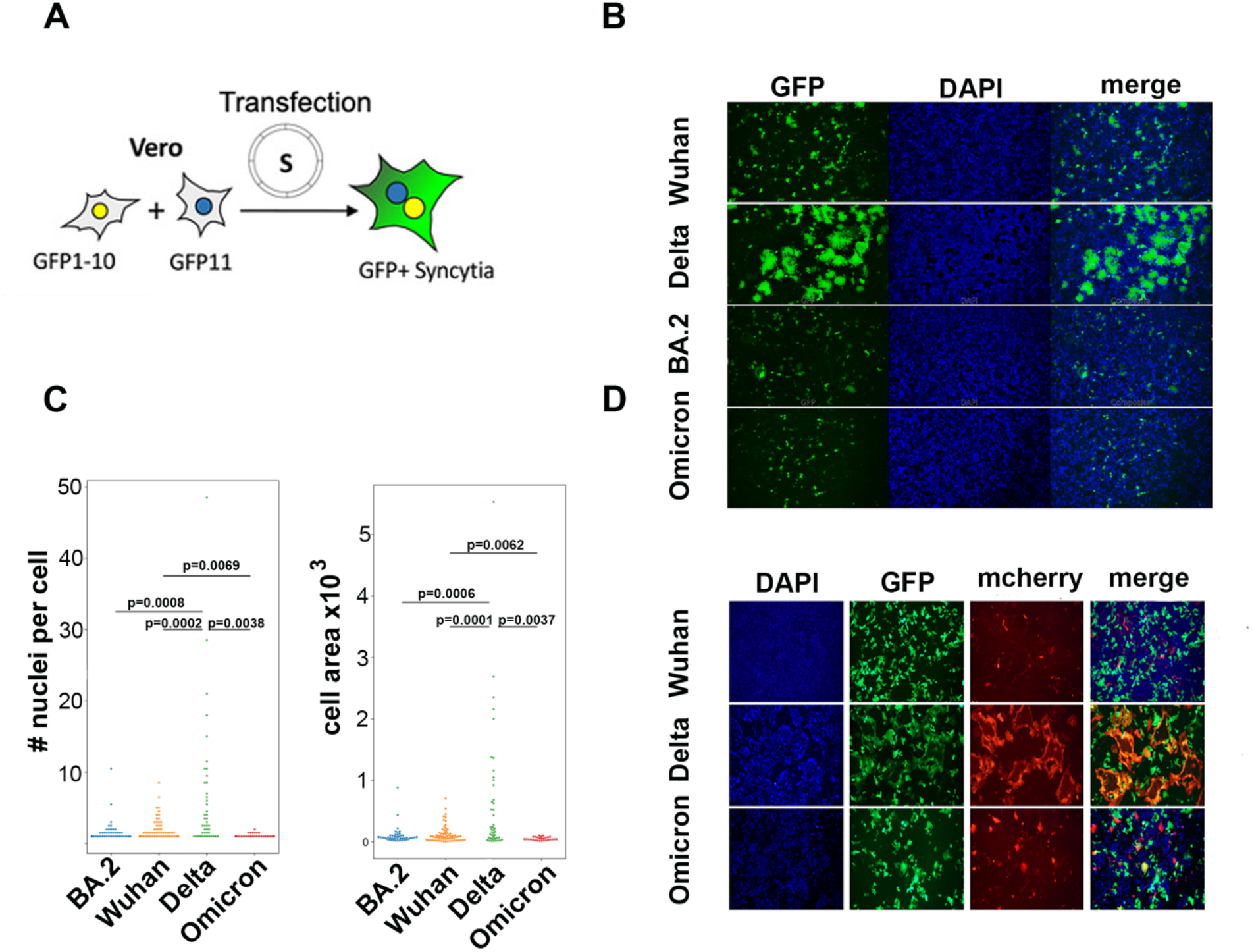
Delta spike promotes enhanced fusogenic activity and syncytia formation relative to the Wuhan-Hu and Omicron SARS-CoV2 spike. **A**. Vero-GFP 10 and 11 represent cells that express each of the GFP subunit. GFP expression is restored only upon cell fusion that is dependent on SARS-CoV-2 spike. **B**. Vero-E6-GFP split were transfected with the indicated spike proteins from Wuhan-Hu, Delta BA.2 and Omicron. 48 h post transfection cells were imaged for their GFP expression as a measurement for cell fusion. **C**. Fusion potential for Wuhan-Hu, Delta and Omicron spike proteins was quantified by imaging GFP and DAPI expression and calculating number of nuclei per cell and cell area as a measurement of fusion. Image analysis and quantitation was calculated based on the expression of GFP and DAPI and was performed using the Scikit-image library [56] in Python 3.1. **D**. Visualization of fusion and syncytia formation - Vero cells stably expression either mcherry or GFP were transfected with the indicated spike. 48 h post transfection cells were imaged for syncytia formation measured by overlapping of mcherry and GFP expression. using Zeiss microscope.

We further imaged cell fusion between Vero cells that stably expressed either mecherry or GFP reporter proteins. Cells were then mixed together at equal numbers and transfected with the indicated spike proteins from either Wuhan-Hu, Delta or Omicron. Syncytia formation, as measured by GFP/mcherry expression overlap per nuclei, was documented using microscopy imaging (**Figure 2D**). We confirmed that the Wuhan-Hu (P681) and Omicron (H681) spike proteins could mediate only a modest cell fusion as shown by their low number of syncytia. On the other hand, Delta spike, carrying the R681 mutation, efficiently enhanced fusion potential and formed syncytia.

### Omicron and Wuhan-Hu exhibit similar plaque formation morphology which is different from Delta upon infection with live viruses

The ability of the Wuhan-Hu SARS CoV-2 or its Delta and Omicron variants to induce cell–cell fusion and syncytium formation was also monitored using live viruses. We envisioned that poor fusion ability will impair viral infection through cell-cell contact. We therefore monitored plaque formation and size by employing plaque assays with live viruses. Following infection, cultures were layered with growth medium containing a semi-solid agarose, which inhibits cell-free infection and viral spread, favoring cell-cell infection. Infected cells were then incubated at 33°C, fixed with 4% PFA and washed before staining with Gentian Violet, which stains cells. Our assay showed that Wuhan-Hu and Omicron viruses exhibited similar plaques morphology, which were different than those of Delta. Delta formed more plaques than Wuhan-Hu and Omicron, which were less unified in their morphology and smaller in their size. These results indicate that in a semi-fluid medium that limits cell-free viral spread, Delta still can form plaques via cell-cell interaction and syncytia formation (**Figure 3**).

**Figure 3:**
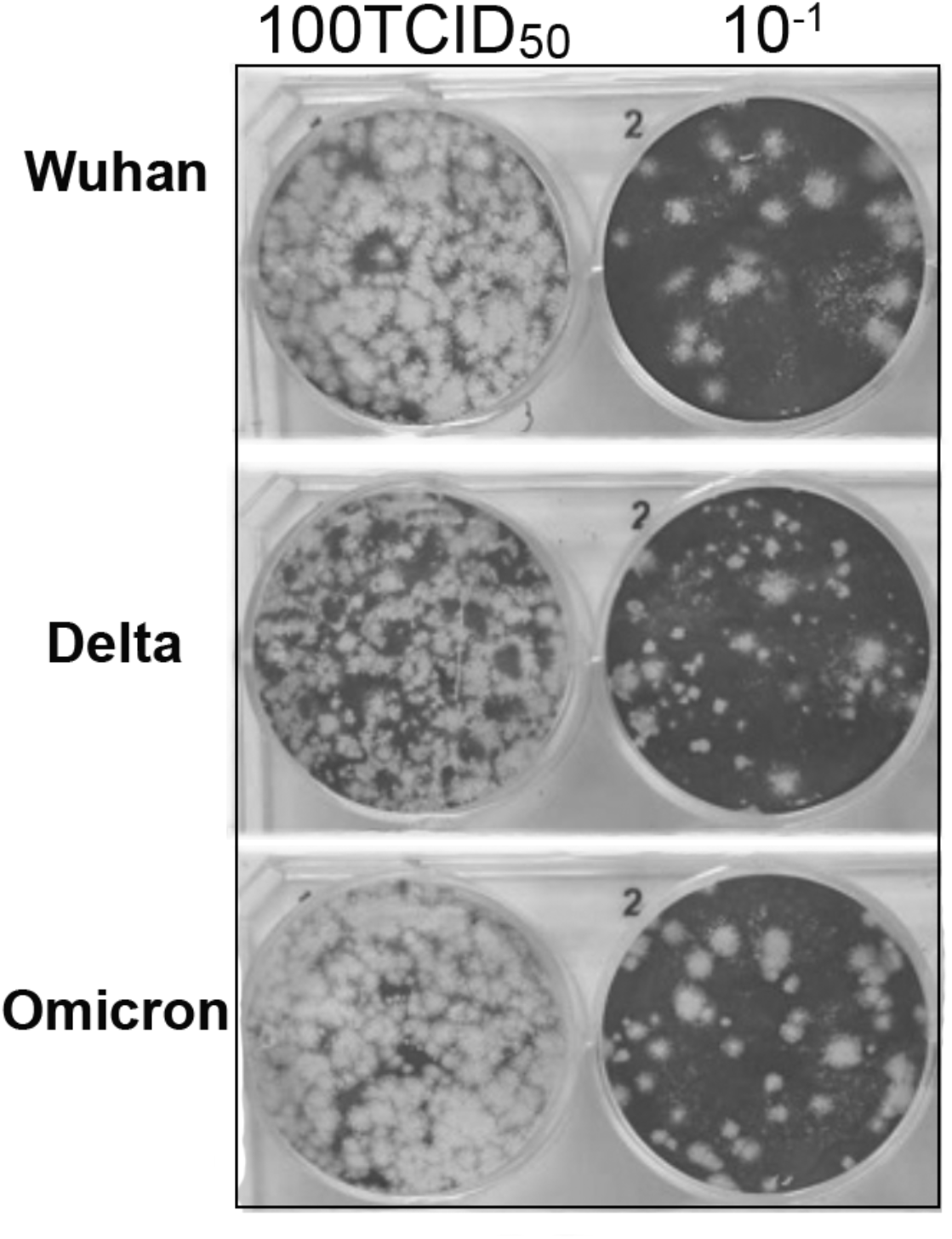
Delta SARS-CoV-2 displays a differential plaque morphology upon infection. Vero E6 cells were seeded in 6 well plates at a concentration of 10^6^ cells/well with 10% FCS MEM-EAGLE medium, and incubated at 37°C. 24 hr later, cells were infected with the following SARS-CoV-2 variants: 1. Wild Type sub lineage B.1.1.50 (hCoV-19/Israel/CVL-45526-ngs/2020), 2. Delta, B.1.617.2 (hCoV-19/Israel/CVL-12804/2021), and 3. Omicron BA.1 (hCoV-19/Israel/CVL-n49814/2021) as 10-fold serial dilutions of 100TCID50 for 1 hour at 33°C. The cells then were washed and MEM-EAGLE medium containing 2% FCS and 0.5% NuSieve agarose was added. Following, cells were incubated at 33°C for 5 additional days. Prior to fixation, 10% Trichloroacetic acid was added for 10 minutes, following by washing of the agarose and staining with Gentian Violet containing 4% Formaldehyde.

### P681 position is critical for fusion and syncytia formation of SARS-CoV-2 spike

Both Delta and Omicron spike proteins carry within their poly-basic _681_PRRAR/SV_687_ motif a mutation at position P681, being either P681H in Omicron, or P681R in Delta. To determine the functional significance of these mutations in enhancing fusion and syncytia formation between SARS CoV-2 target cells, we mutated the P681 residue of the Wuhan-Hu spike to R681 that is displayed on the spike of Delta. Insertion of a single R681P mutation in the context of Wuhan-Hu spike and expression in Vero-GFP-split cells, resulted in an enhanced GFP expression, indicating that fusion and syncytia formation potentials were restored to the levels exhibited for Delta spike (**Figure 4A**). Conversely, switching R681 in Delta spike to the Wuhan-Hu version - P681, efficiently abolished enhanced fusion potency and formed syncytia that were low in their numbers and small in their size (**Figure 4A**). We then mutated the spike of Omicron H681 into R681 and expressed these spike proteins in Vero-GFP-split cells. We document an enhanced fusion potential and increased syncytia formation upon mutating the Omicron H681 to R681 spike (**Figure 4A)**. We further quantitated the levels of GFP expression, as a measure for fusogenic potential and syncytia formation (**Figure 4B**) and concluded that P681 in the polybasic region of spike is critical for fusion activity and for syncytia formation.

**Figure 4:**
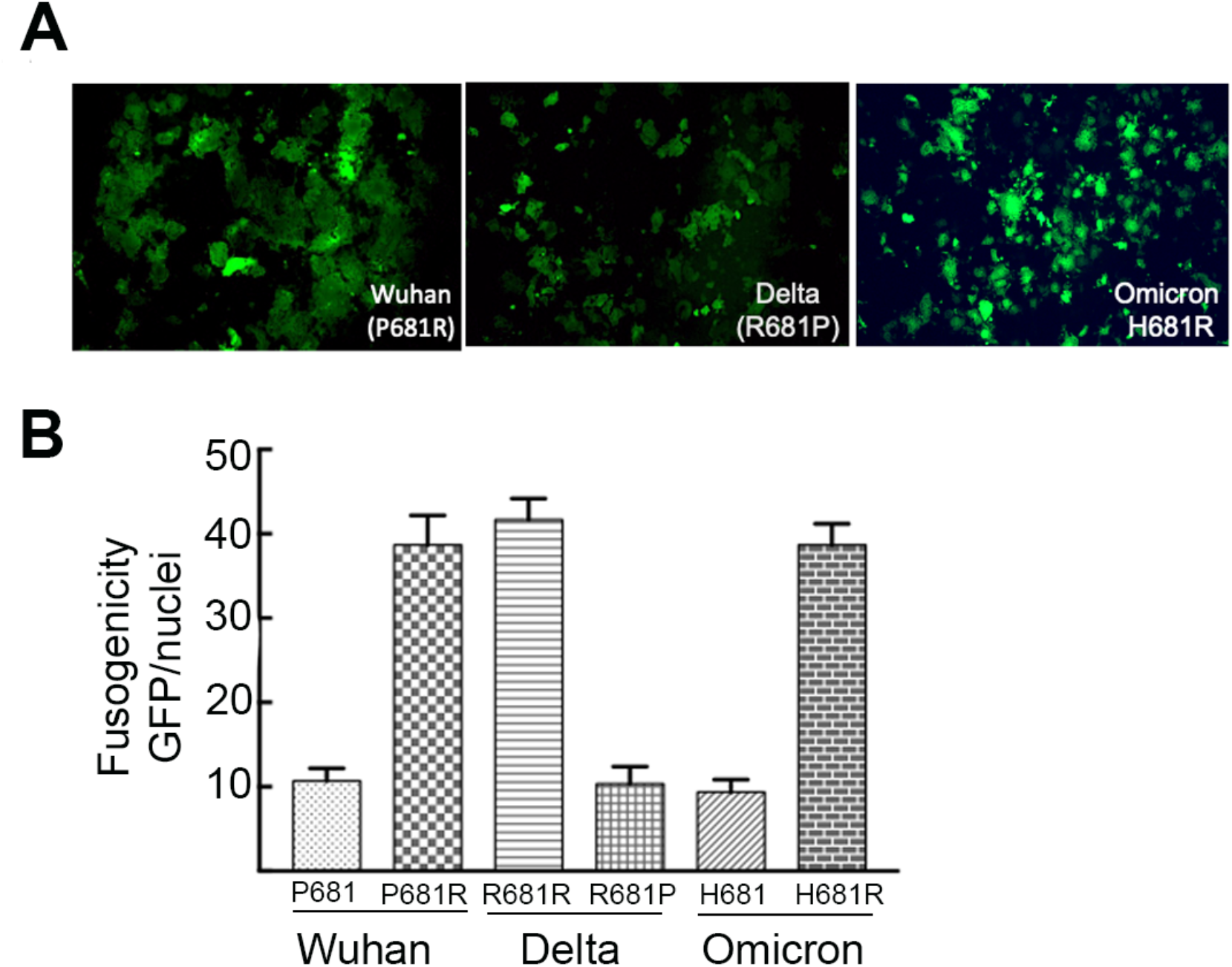
P681 dictates cell fusion and syncytia formation of SARS-CoV-2 spike. **A**. Equal numbers of Vero-E6 cells expressing either GFP1-10 or GFP 11 (Vero GFP-split) were mixed and then transfected with the indicated Wuhan-Hu, Delta or Omicron spikes spike proteins carrying the specified point mutation within the P618 residue of the poly basic motif. 72 hr post transfection cells were monitored for their GFP expression by microscopy. **B**. Quantitation of fusogenicity using image J (GFP area/number of nuclei).

## Discussion

Syncytia formation has been described as a hallmark of many viruses including SARS-CoV-2 [60, 61]. Entry of SARS CoV-2 and subsequent infection are mediated by the interactions between the spike expressed on the cell surface of infected cells, and the human ACE2 receptor [62]. Furthermore, cell fusion and formation of syncytia in the lung of infected patients, correlate with clinical symptoms, severe disease and patient mortality [61, 63, 64]. Therefore, characterizing the residues/mutations in spike that are involved in cell fusion and syncytia formation is critical.

Along RBD mutations within spike that have been extensively studied for their effects on viral infectivity and vaccine-induced neutralization sensitivity, circulating Alpha, Beta and Delta variants also harbor mutations within their Furin Cleavage site (FCS), primarily at the P681 spike residue. In this study, we used pseudoviruses and monitored the neutralization potential of a cohort of sera drawn from fully vaccinated individuals – four months after third dose against the Wuhan-Hu SARS CoV-2 and its Delta and Omicron variants (**Figure 1**). We show that, relative to wild-type Wuhan-Hu pseudovirus, the efficiency of our samples to neutralize Omicron was substantially reduced – x26 fold. Furthermore, a decrease in the tested sera to neutralize the Delta pseudovirus was also reduced x13 fold relative to the Wuhan-Hu SARS CoV-2. We also demonstrated that one of the main drivers for this neutralization decrease was an enhanced infectivity level of Omicron, which was x5 fold higher than the Wuhan-Hu pseudovirus (**Figure 1**). Overall, we can conclude that over time, the efficiency of the vaccine-elicited immune response is waned, rising major concerns regarding the protection this vaccine provides against the Omicron variant and the need for an additional dose.

We further investigated the effects of mutations at position 681 polybasic region within the FCS that appear in different circulating variants on fusion potential and syncytia formation of cells expressing the spike protein. When expressed in Vero target cells, our data show that Wuhan-Hu and Omicron spike proteins promote modest levels of fusion potential and syncytia formation (**Figure 2**). In contrast, the spike of the Delta efficiently promoted enhanced cell fusion and generated large syncytia between target cells (**Figure 2**). Our transit expression experiments were also confirmed with live viruses, demonstrating that Delta forms different size plaques which are higher in numbers when compared to Omicron and Wuhan-Hu (**Figure 3**). Furthermore, the P681 residue has been mapped as critical for the fusogenic activity of the different spike proteins. Changing P681 in the Wuhan-Hu spike into R681 seen in Delta, restored fusogenicity and syncytia formation. Similarly changing R681 of the Delta spike into P681 or H681 positioned within spike of Wuhan-Hu or Omicron, abolished the fusion phenotype seen in Delta spike (**Figure 4**).

The polybasic PRRAR motif is unique in the spike of SARS CoV-2. However, its functional significance for viral infection is still debatable. Recent work has confirmed that this region is important for viral transmission as it provides an evolutionary advantage for the virus to enter its target cells at the cell surface by promoting membrane fusion between the virus and the cell and between infected cells [49, 50, 65, 66]. Such syncytia potentially facilitate viral replication, dissemination and immune evasion, causing cytopathic effects and wider tissue damage. In this scenario, the virus successfully evades the innate IFITM2 response upon entry and efficiently spreads [45, 51, 67]. On the other hand, in viruses that lack the polybasic PRRAR-region, viral entry is mediated through endosomes, and therefore exposed to the IFITM2-innate anti-viral response. In Vero E6 cells, that do not express the TMPRSS2 protease and lack of innate response, SARS-CoV-2 deleted FCS gain an advantage, potentially because there is an increase in its spike stability, and premature shedding of the S1 subunit that abrogates receptor binding [68]. Knockout of TMPRSS2 or deletion of the poly-basic sequences, allows SARS-CoV-2 to mediate viral entry in a pH-independent manner, in part to mitigate against IFITM-mediated restriction and promote replication and transmission [51]. FCS deletion also results in impaired infection, which is due to low viral titers that are shed from an infected ferrets animal model, resulting in reduced transmission to cohoused sentinel animals [45]. This report also showed that the TMPRSS2-mediated entry is more efficient in viruses that express the FCS polybasic region [45]. Moreover, given the fact that upon propagating the virus in cell culture, FCS sequence is lost or mutated, while in infectious viral isolates these mutations surface at a very low level, further strengthen the assumption that FCS sequence is critical for viral transmission, only in clinically relevant systems and infection by live virus [45]. In another recent work, the importance of the polybasic FCS region was further reinforced. FCS-P681R mutation within Kappa-SARS-CoV-2 augmented syncytium formation, thus contributing to increased infectivity of the Alpha and Beta and SA variants [46]. Other residues that showed to be involved in cell fusion included D614G, K417N and to a lesser extent E484K. However, in our study we could not see such an effect on viral infection nor neutralization potential presumably since our infectivity assays employed single round pseudoviruses and focused only on the early step of entry and infection, with no ability to extend the effects on late stages on virus release of infectious particles. Moreover, we did not perform our assays in a clinically relevant system, where viral infection is dependent on TMPRSS2 and requires membrane fusion, thus facilitating syncytium formation. Other studies have also demonstrated that the Delta variant exhibits higher fusion potential relative to the wild-type SARS CoV-2 or the Alpha - P681H mutation [52, 69-71]. Moreover, Omicron replicates more slowly than Delta upon over expressing TMPRSS2. Use of specific inhibitors targeting the endocytic or TMPRSS2 entry pathways further concluded that while Omicron variant uses mainly the endocytic pathway for entry target cells, Delta uses both the endocytic and TMPRSS2 pathways. The difference in entry pathway between Omicron and Delta variants may have an implication on the clinical manifestations or disease severity [52]. Our study confirms these data and further define the P681 residue within FCS region of spike as the key residue that dictates cell-cell fusion and syncytia formation.

## Acknowledgements

This work was supported by the Israeli Ministry of Science and Technology for RT (MOST; grant #3-16897), the Israel Science Foundation for RT (ISF; Research Grant Application no. 755/17) and the Ben-Gurion University of the Negev COVID-19 Research Task Force for RT.

The work of BR was supported by ISF grants numbers: 1416/19 and 2841/19.

## Author Contribution

A.K. and R.T. conceived the study and analyzed the data. A.K., DK and ILC performed experiments and analyzed the data. N.A analyzed the effects of live virus on syncytia formation. A.O and B.R performed analysis on the images of Vero split cells. E.R helped in obtaining sera samples and submission the request to obtain human sera samples from the Institutional Helsinki Review Board. R.T. wrote the manuscript.

## Declaration of Interest

The authors have no conflicts of interest to declare.

